# P*Can*PIE: A group I intron platform for efficient circRNA synthesis at ambient temperatures

**DOI:** 10.64898/2026.05.11.724386

**Authors:** Ruben Warkentin, Anna Marie Pyle

## Abstract

Ribozyme-based permuted intron-exon (PIE) systems offer a protein-independent route to circRNA production, but existing platforms require elevated temperatures that promote RNA degradation. Here we report the first application of the *Candida albicans* mitochondrial large subunit (*C.a*.mtLSU) group I intron as a PIE platform for circRNA synthesis, which we term P*Can*PIE (Pyle lab *Candida* PIE). We evaluated three peripheral stems, P5, P6b, and P8, as permutation sites and demonstrated that all three support circularization under near-physiological conditions (25°C, 6 mM MgCl_2_), without the 55°C heating step required by existing PIE systems. Kinetic analysis revealed that permutation site does not affect the observed splicing rate constant but does influence P*Can*PIE folding and therefore influences circularization efficiency. The P6b permutation yielded the highest circularization efficiency, with 95 % of the precursor splicing to produce circRNA. Optimization of spacer sequences flanking the circRNA payload eliminated interference from structured native exon sequences and enabled efficient circularization of RNAs up to 1,657 nt, including structured, repetitive, and naturally occurring sequences. Together, these results establish P*Can*PIE as a versatile and near-physiologically active addition to the group I intron PIE toolkit.

## INTRODUCTION

Circular RNAs (circRNAs) have emerged as an important topological class of RNAs, exhibiting greater stability than linear RNAs due to their resistance to exonuclease degradation. CircRNAs carry out diverse biological functions, including miRNA sponging, protein scaffolding, and translation (Chen 2020). Together, these properties have motivated growing interest in their use for biotechnological and therapeutic applications.

Several strategies exist for the synthesis of circRNAs in vitro, including chemical ligation, enzymatic ligation, and ribozyme-based approaches, but each have their drawbacks. Recent advances in chemical ligation enable synthesis of gene-sized circRNAs; however, efficiencies for production of large circRNAs (4000 nt) are low and intermolecularly ligated byproducts are produced (Wasinska-Kalwa et al. 2025). Furthermore, chemical ligation requires non-standard modified RNA termini to produce the junction. While protein-based enzymatic reactions are routinely used to produce intermolecular ligated products, RNA circularization efficiency by protein enzymes is highly inefficient (Kim et al. 2023). By contrast, ribozyme-based approaches can improve the yield of circRNA products and avoid the accumulation of intermolecular spliced by-products, making them an attractive alternative for in vitro circRNA production.

Many ribozyme-based circRNA synthesis methods use permuted intron-exons (PIEs) that are derived from split group I introns flanking an intervening RNA payload of interest (**Figure 1**). Self-splicing of these “circularly permuted” RNA precursors releases the payload sequence as a covalently closed circle. The PIE approach was first demonstrated by Puttaraju and Been (1992), who showed that the *Anabaena* and *Tetrahymena* group I introns can be permuted at multiple intron positions, yielding a cyclic product. Despite these early reports of flexible permutation site positioning, subsequent work in PIE systems has focused almost exclusively on constructs that have been permuted by splitting the P6 region (Wesselhoeft et al. 2019; Chen et al. 2023; Du et al. 2024). To our knowledge, there has been no optimization of permutation site selection and no investigations on the effect of permutation position on intron folding and circularization efficiency. Additionally, all described PIE systems require high temperatures (55°C) to splice efficiently, which could be explained by PIE systems that are kinetically trapped in misfolded conformers that must be resolved before the active structure can form. Recently, an alternative ribozyme-based strategy called “complete self-splicing intron for RNA circularization (CIRC)” partially addresses this issue. CIRC is based on a P6 permuted *Anabaena* intron, but places the entire permuted intron upstream of the exon rather than flanking it, skipping the first transesterification reaction (Shen et al. 2025). This approach enables splicing of large RNAs up to 14 kb, but still requires 55°C for efficient circularization (Shen et al. 2025). These elevated temperatures promote RNA degradation and limit the utility of these systems for efficient production of large circRNAs.

**Figure 1.**
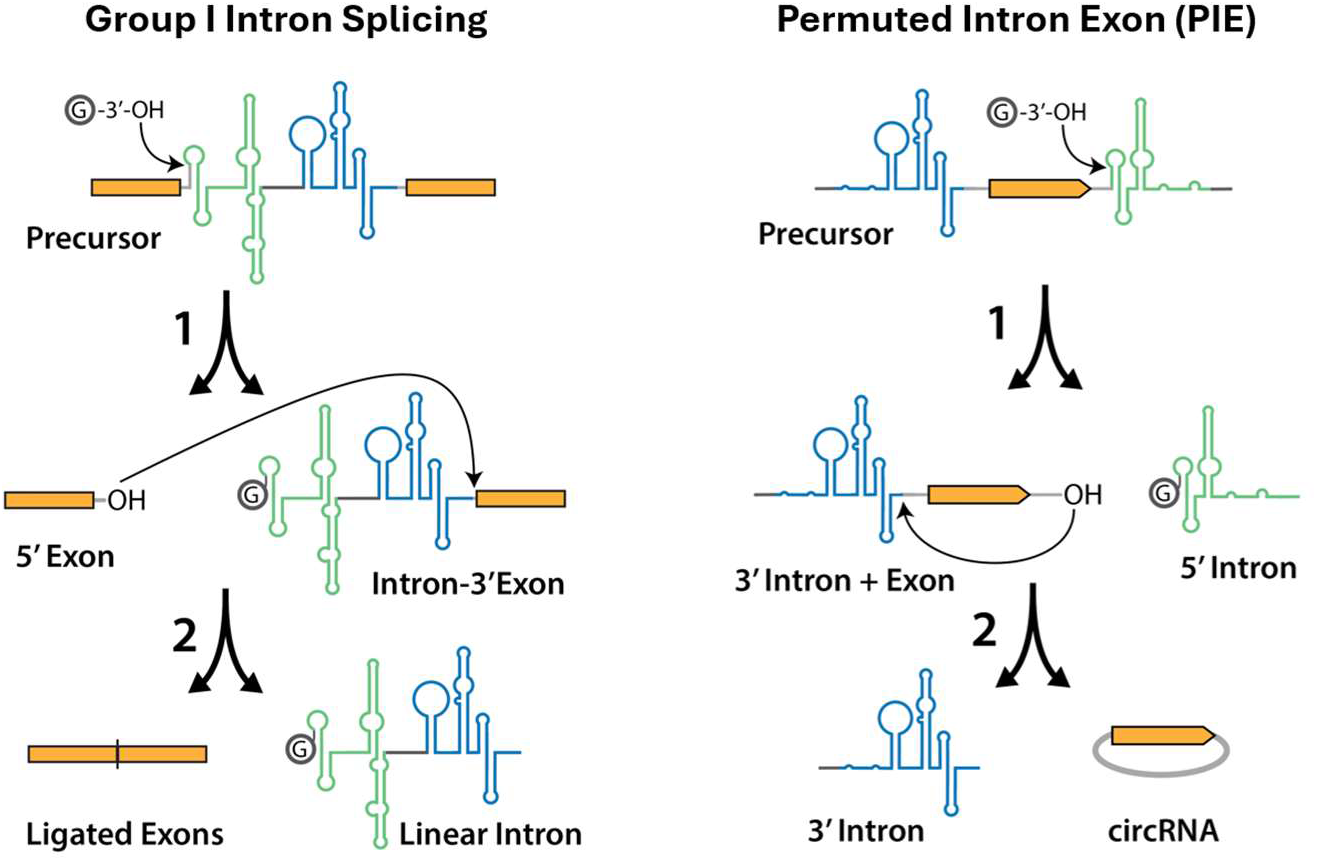
Schematic of group I intron and permuted intron-exon (PIE) splicing. (**Left**) Canonical group I intron splicing proceeds via two transesterification reactions. In the first step, an exogenous guanosine attacks the 5′ splice site. In the second step, the free 3′-OH of the 5′ exon attacks the 3′ splice site, ligating the two exons and releasing the excised intron. (**Right**) In the PIE strategy, the intron is circularly permuted such that the 3′ half-intron is placed upstream and the 5′ half-intron downstream of the sequence to be circularized. The same two-step transesterification chemistry joins the ends of the intervening sequence producing circular RNA (circRNA). Exons are shown in yellow; intron sequences in green and blue.

The *Candida albicans* mitochondrial large subunit group IA1 intron (*C.a*.mtLSU GpIA1) presents a compelling alternative intron for PIE engineering. We recently showed that the *C.a*.mtLSU GpIA1 intron reacts rapidly and with high efficiency under near physiological conditions (25 °C and 3 mM MgCl_2_) (Liu and Pyle 2024). A subsequent structure determination of *C.a*.mtLSU GpIA1 from our laboratory revealed a distinctive catalytic core stabilized by a P7ext-P9.1 tertiary interaction and an unusual P7ext-P6b pseudoknot (P11) that is characteristic of group IA1 (Liu et al. 2025). Importantly, this intron lacks the alternative P3 pairing that causes kinetic trapping in the *Tetrahymena* intron (Bonilla et al. 2022), which is one of the most commonly-used introns for circRNA synthesis. Together, the reinforced catalytic core and rapid, low temperature splicing suggested to us that this intron might be more tolerant to the structural perturbations imposed by the PIE circRNA synthesis.

In this work, we present the first application and characterization of the *C.a*.mtLSU GpIA1 as a PIE platform for circRNA synthesis, which we term Pyle lab *Candida* PIE (P*Can*PIE). We compare multiple stems as permutation sites, finding that permutation site choice does not affect splicing rate constants but does influence intron folding efficiency and subsequently the circularization efficiency. In addition, we optimized the spacer sequences flanking the circRNA payload, enabling efficient circularization of RNAs up to 1.6 kb. Together, these studies establish P*Can*PIE as the first PIE platform capable of efficient circRNA synthesis at ambient temperatures.

## RESULTS

### Structure-guided permutation design of the P*Can*PIE

To develop a PIE system capable of synthesizing circRNA efficiently, we identified structurally permissive regions of the highly efficient *C.a*.mtLSU GpIA1. Analysis of the cryo-EM structure (PDB: 9MQS) revealed three peripheral stems, P5, P6b, and P8, that are spatially separated from the catalytic core and minimally involved in long-range tertiary contacts (**Fig. 2A**). Their respective positions suggested that each site could accommodate the backbone discontinuity introduced by circular permutation without disrupting global folding or active-site architecture. Guided by this rationale, we generated PIE constructs by repositioning the 5’ and 3’ ends of the intron to flanking sides of each stem, introducing short sequence adjustments to maintain base pairing (**Fig. 2C**).

**Figure 2.**
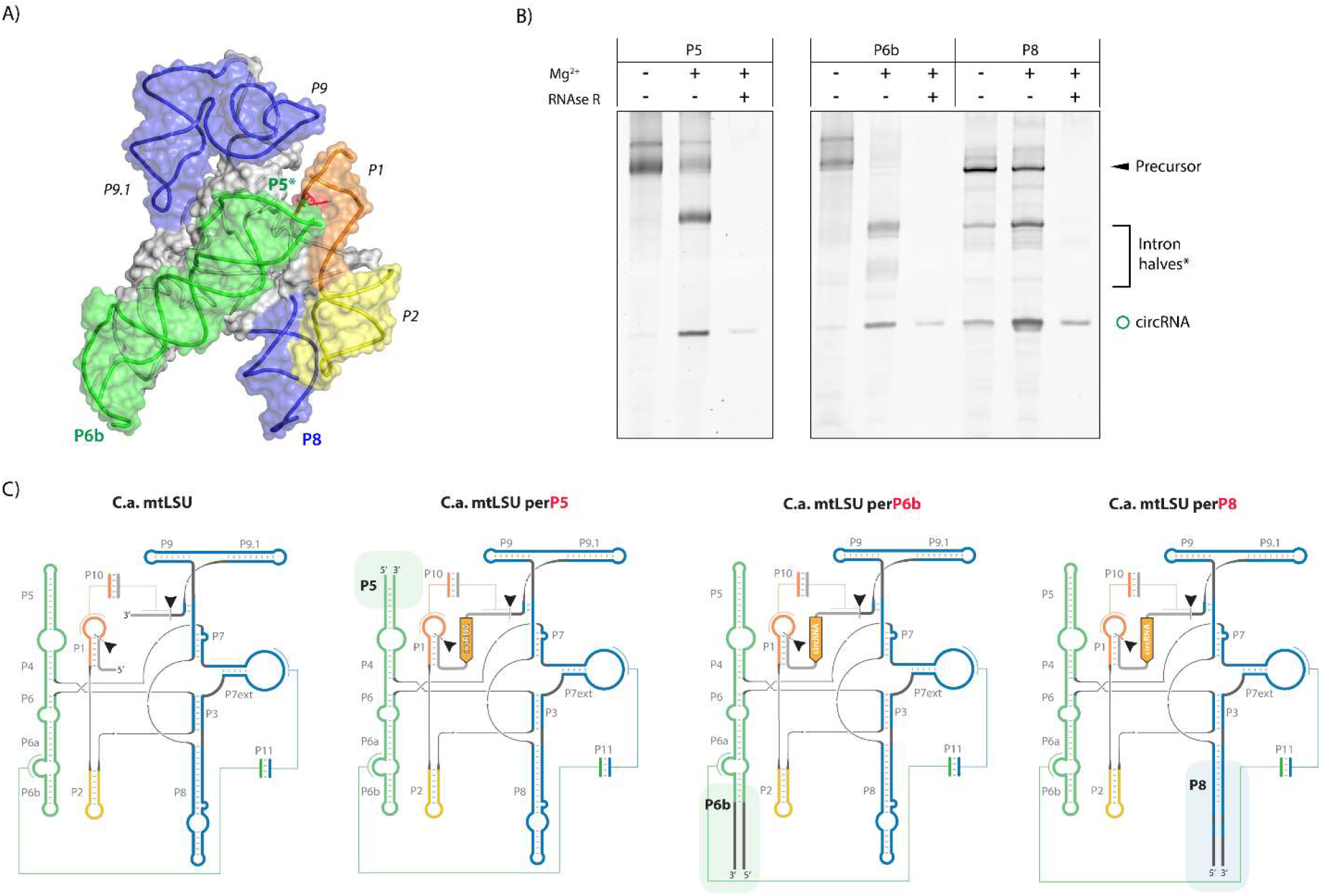
Structure-guided permutation design of *C.a*.mtLSU GpIA1 and circRNA synthesis. (A) Secondary structure diagram of the *C.a*.mtLSU GpIA1 intron highlighting peripheral stems P5, P6b, and P8 as candidate permutation sites. (B) Denaturing PAGE analysis of in vitro splicing assays for P5, P6b, and P8 PIE constructs, with and without RNase R digestion, confirming circRNA production. (C) Schematic of PIE construct design showing repositioning of intron ends to each permutation site with sequence adjustments to maintain local base pairing.

We then tested whether these structure-guided permutations support self-splicing and circularization. In vitro splicing assays revealed that permutations at P5, P6b, and P8 each yielded an RNA species corresponding to the expected circRNA product that is resistant to RNase R digestion (**Fig. 2B**) (Suzuki et al. 2006). These results demonstrate that multiple peripheral stems of the *C. albicans* intron can be used as functional permutation sites for circRNA synthesis. We next compared the splicing efficiencies of the P5, P6b, and P8 variants and assessed whether the redesigned intron retained its ability to splice under near-physiological conditions.

### Permutation site affects intron folding but not splicing kinetics

To determine whether permutation site influences catalytic performance, we compared the splicing efficiencies of the P5, P6b, and P8 P*Can*PIE constructs using radiolabeled precursor RNA in time-course assays under near-physiological conditions (25°C, 6 mM MgCl_2_). All three variants produced the expected circRNA product with nearly identical observed rate constants (*k*_obs_) of 0.15 ± 0.02 to 0.17 ± 0.01 min^−1^ (**Fig. 3**). This indicates that none of the permutation sites that we tested significantly affect the intrinsic catalytic rate of the intron. However, despite their similar *k*_obs_ values, the three permutations differed substantially in their reaction amplitudes (extent of reaction). The P6b permutation had highest amplitude, with only 5.3 ± 0.1% precursor remaining at reaction completion (94.7 % circularization efficiency), which is more than 3-fold higher than observed for the P5 and P8 constructs. This difference in reaction amplitude indicates that the permutation site can influence the fraction of molecules that fold efficiently into a splicing-competent conformation. Based on these results, we selected the P6b permutation as the starting design for generating larger circRNAs.

**Figure 3.**
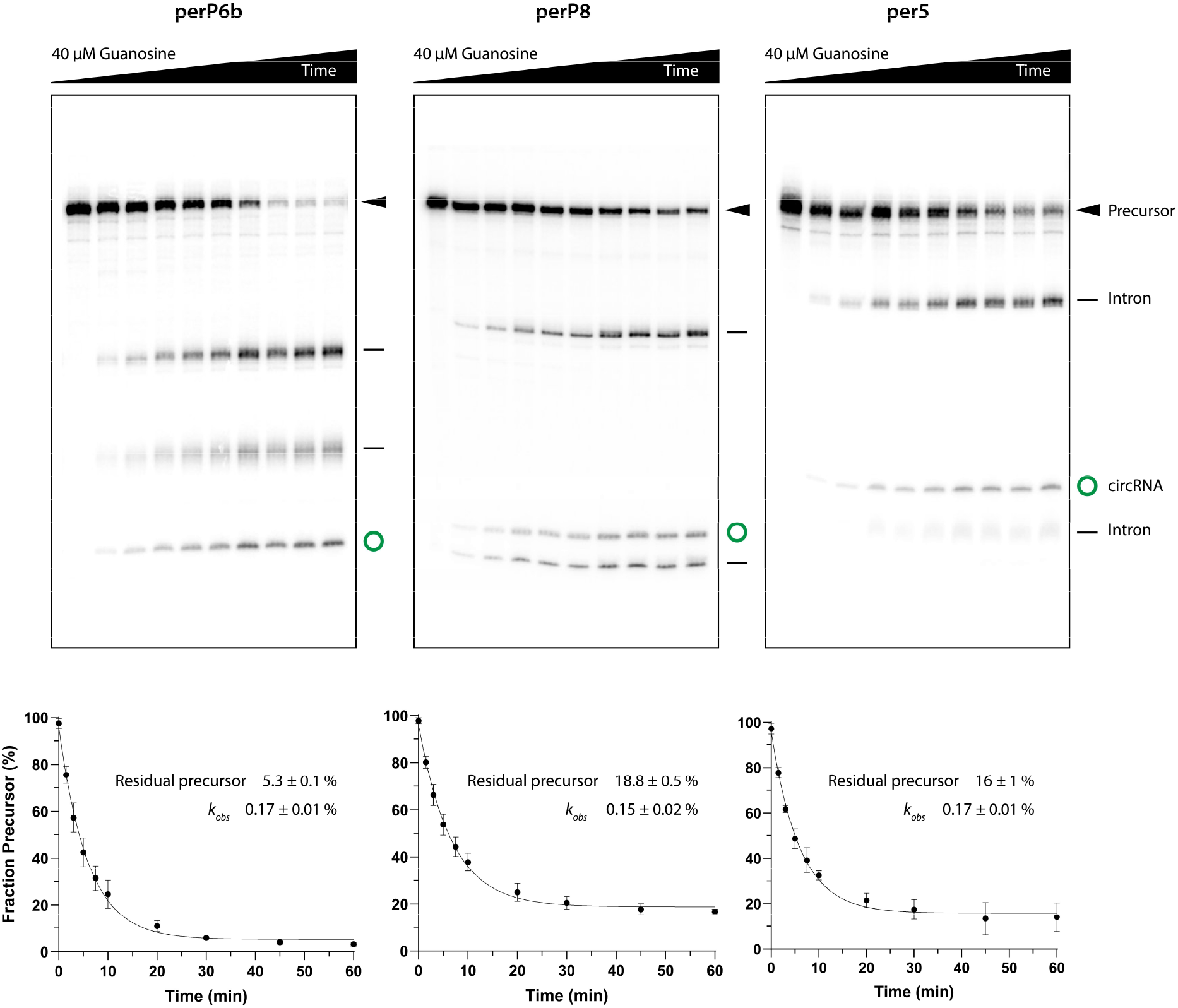
Kinetic analysis of P5, P6b, and P8 P*Can*PIE constructs shows efficient splicing at low temperatures and magnesium concentrations. Time-course splicing assays using radiolabeled precursor RNA at 25°C, 6 mM MgCl_2_ were analyzed by 5 % polyacrylamide/8M urea gel. Plots show fraction of precursor remaining over time fit to a single exponential decay. Inset table shows *k*_obs_ and amplitude values (mean ± SD, n = 3) for each permutation site.

### Spacer optimization enables circularization of larger RNAs

Having established P6b as the most robust permutation site, we next tested whether this design could support synthesis of larger circRNAs. Two constructs containing 200 nt payloads were generated: one containing an extended version of the native *C.a*.mtLSU GpIA1 exon sequence, and another containing the original 103 nt sequence supplemented with 97 nt of unstructured actin sequence (circActin). Splicing of the extended mtLSU construct failed, producing aberrant products (**Fig. 4D**), while the circActin construct showed only partial activity with substantial precursor remaining. These results indicated that simply increasing circRNA size can interfere unpredictably with productive intron folding.

**Figure 4.**
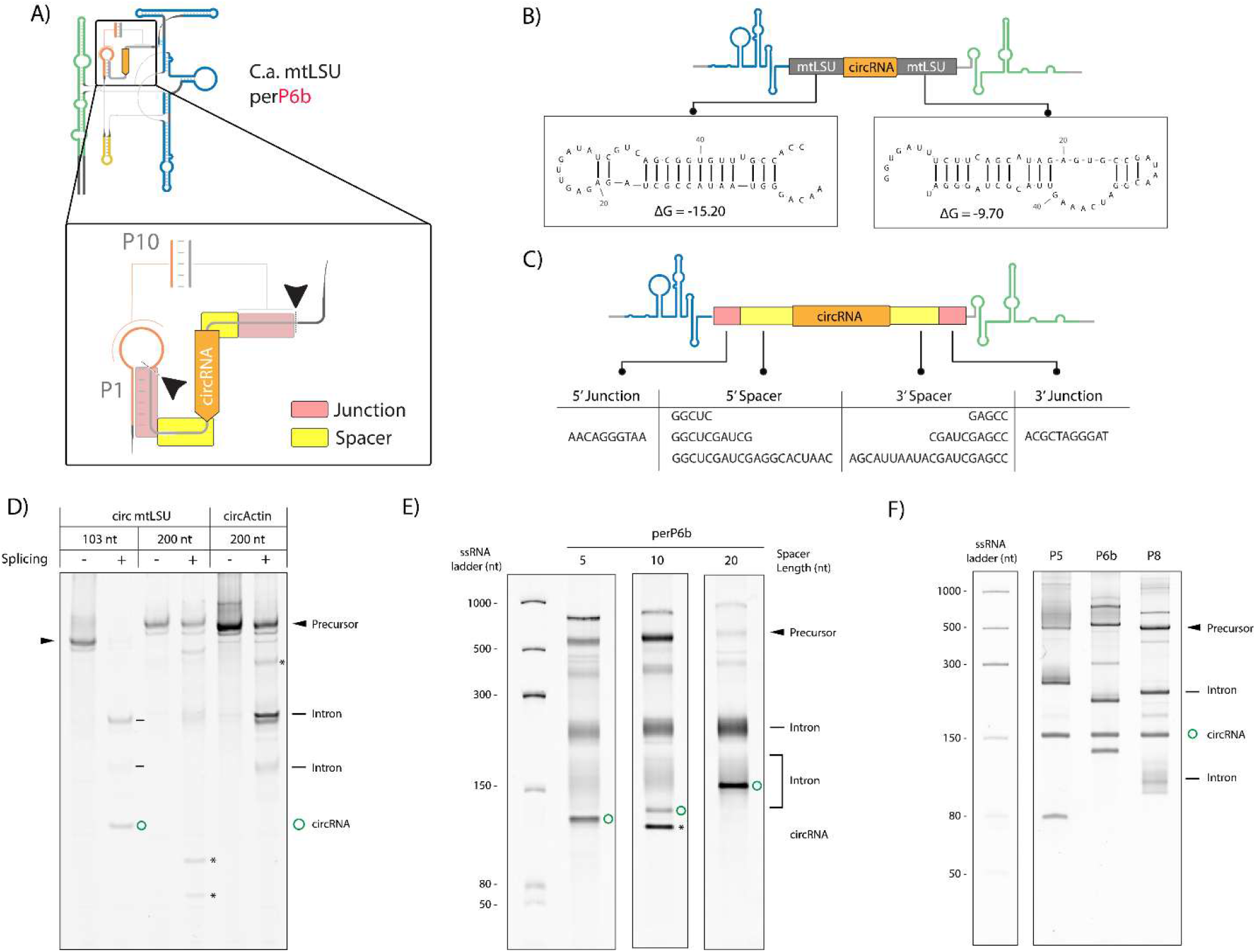
PIE spacer optimization for efficient circularization. (A) Schematic showing the invariant P1/P10 junction sequences and the adjacent spacer regions that were redesigned. (B) Secondary structure predictions of native mtLSU exon sequences showing stems predicted to interfere with intron folding. (C) Redesigned PIE construct with neutral spacer sequences and 3×FLAG circRNA payload. (D) Denaturing PAGE showing failed splicing of the extended mtLSU construct. (E) Comparison of 5, 10, and 20 nt spacer designs showing circRNA production. (F) Denaturing PAGE confirming that the 20 nt spacer design is compatible with P5, P6b, and P8 permutation sites.

The *C.a*.mtLSU GpIA1 intron contains rRNA-derived sequences, leading us to hypothesize that structured elements within exons, and particularly the mtLSU exon region, may form unintended interactions with the intron or circRNA and disrupt folding. Secondary structure predictions supported this idea, showing that mtLSU sequences flanking the circRNA region form stems that may interfere with intron architecture (**Fig. 4B**). To prevent these interactions, we redesigned the junction and spacer regions. As shown in Figure **4A**, junctions corresponding to the essential P1 and P10 pairing regions cannot be altered, although the adjacent spacer sequences do not participate in intron structure and can be readily modified. We replaced the mtLSU-derived spacer with a neutral sequence and used a 3×FLAG tag as the circRNA payload, as in other PIE systems (**Fig. 4C**) (Du et al. 2024). Spacer lengths of 5, 10, and 20 nt were tested. All designs produced circular RNA, with circRNA size increasing proportionally with spacer length (**Fig. 4E**). The 10-nt spacer showed evidence of mis-splicing, whereas the 20-nt spacer yielded the strongest circRNA band and minimal precursor, indicating that the 20-nt spacer is the most effective design.

We then tested whether this optimized spacer design is compatible with the P5 and P8 permutation sites. All permuted designs produced the expected circRNA band (**Fig. 4F**), with varying levels of efficiency. Together, these results show that removing the structured exonic elements and introducing optimized spacers can restore efficient splicing and that a generalized spacer design can be used for different permutation sites. We next tested whether these designs can be used to generate significantly larger circRNAs.

### P*Can*PIE P6b efficiently produces large and sequence-diverse circRNAs

The ability to generate large circRNAs is an important benchmark for evaluating the versatility of PIE systems, as many biologically relevant circRNAs require regulatory RNA elements and coding sequences, often exceeding 1,000 nt. These features impose substantial folding and steric constraints on the precursor RNA, reducing the splicing efficiency of most existing PIE platforms or requiring elevated temperatures for splicing of larger payloads.

To assess the limits of the P*Can*PIE platform, we selected a panel of circRNAs that increase in size and structural complexity. These included a 250 nt circActin construct representing an unstructured RNA, a 420 nt circATV construct containing internal repeats and predicted secondary structure, a 934 nt circZnf609 corresponding to a naturally occurring circRNA, and a 1,657 nt circCVB3 EGFP construct that incorporates both a coding region and a structured IRES element. All four designs produced circular RNA during in vitro transcription, demonstrating that the redesigned junction and spacers enable efficient circularization across a broad range of sequence lengths and structural contexts (**Fig. 5A, B**). The successful circularization of the 1.6 kb coding construct demonstrates that P*Can*PIE can tolerate the structural and steric demands imposed by long ORFs and IRES-containing sequences. The identity of each circRNA was confirmed by RT-PCR and nanopore sequencing (**Fig. 5C**). Together, these data indicate that the redesigned junction and spacer support robust circularization of a wide variety of circRNA sequences, including long, structured, and naturally occurring RNAs, without requiring additional optimization. This establishes a flexible and broadly applicable platform for generating large and diverse circRNAs.

**Figure 5.**
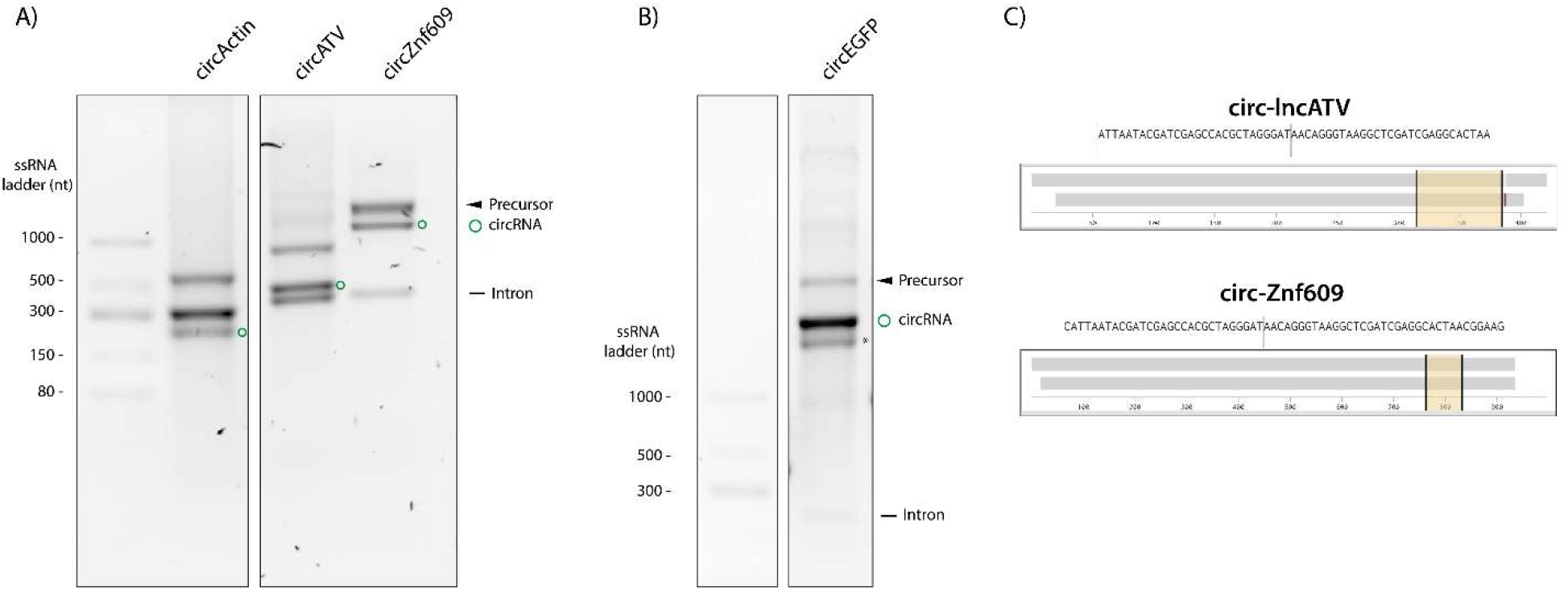
P*Can*PIE P6b efficiently produces large and sequence-diverse circRNAs. (A) Schematic of circRNA constructs of increasing size: circActin (250 nt), circATV (420 nt), circZnf609 (934 nt), and circCVB3-EGFP (1,657 nt). (B) Agarose gel electrophoresis of circRNA products with and without RNase R treatment. (C) RT-PCR analysis using divergent primers and nanopore sequencing confirmation of the back-splice junction for each circRNA construct.

## DISCUSSION

### Permutation site shapes folding efficiency rather than catalytic rate

This study demonstrates that group I introns, and specifically the *C.a*.mtLSU GpIA1, can be permuted at multiple peripheral stems without loss of catalytic activity, and establishes that permutation site choice has distinct effects on intron folding and circularization efficiency. Permutation at P5 and P8 does not impair catalysis per se, but increases the proportion of precursor RNAs that misfold into inactive conformers, resulting in 81-84 % conversion of precursor RNA to circRNA, compared to 95 % for the P6b permutation (**Fig. 3**).

In the case of the *C.a*. mtLSU intron, superior folding efficiency of introns permuted at position P6b may reflect the structural role of P6b in forming the P11 pseudoknot with P7ext, which reinforces the catalytic core of *C.a*.mtLSU GpIA1. Disrupting this peripheral stem through permutation may be particularly well-tolerated because the P11 tertiary interaction buttresses the core and may ensure the two intron halves come together. Permutation at P5 or P8 may impose greater structural complexities on the folding landscape. Importantly, our findings demonstrate that not all group I based PIE systems require high temperatures (55 °C) for splicing, as P*Can*PIE achieves near-complete precursor conversion (95 %) to circRNA at 25 °C (**Fig. 3**).

It is notable that Puttaraju and Been, chose to permute at P5, P6, and P8 in both the *Anabaena* pre-tRNA and *Tetrahymena* pre-rRNA group I introns (Puttaraju and Been 1992). These are the same peripheral stems we selected here on structural grounds, suggesting that these sites are permutation-tolerant across group I intron architectures. More broadly, the tolerance of peripheral loops for large insertions without disruption of the catalytic core is well established in group I intron biology: homing endonuclease ORF families are documented as insertions in peripheral stems including P1, P2, P6, P8, and P9 across diverse intron families and subclasses (Hedberg and Johansen 2013; Haugen et al. 2005). *C.a*.mtLSU GpIA1 carries no ORF in any of its peripheral loops, leaving these stems structurally unencumbered. We speculate that the absence of ORFs permitted *C.a*.mtLSU GpIA1 to evolve a structure optimally tuned for productive folding, unconstrained by the coding sequence compatibility that selectively shapes ORF-harboring introns (Haugen et al. 2005).

### Structured native exon sequences limit circRNA size and spacer design as a solution

The failure of extended mtLSU exon sequences to support efficient circularization highlights an important design consideration for PIE systems derived from introns that are naturally embedded in structured RNA contexts (**Fig. 4**). *C.a*.mtLSU GpIA1 resides within the mitochondrial rRNA, and the flanking exon sequences carry secondary structure that may compete with essential intron contacts when incorporated into a PIE construct. Replacing these sequences with spacers restored efficient splicing and enabled circularization of diverse payloads without payload-specific optimization (**Fig. 4**,**5**). The requirement for a minimum spacer length of 20 nt suggests that sufficient physical separation between the intron termini and the circRNA payload is needed to prevent steric or structural interference at the splice junctions. Similar requirements have been reported for the *Anabaena* and T4 td PIE systems (Wesselhoeft et al. 2019; Du et al. 2024; Shen et al. 2025).

The compatibility of the optimized spacer design with all three permutation sites indicates that the junction architecture is a general solution rather than one tailored to a specific permutation. This modularity is valuable in practice: it means that for the P*Can*PIE system, the same spacer sequences can be used regardless of which peripheral stem is selected as the permutation site, simplifying construct design when applying this platform to new circRNA sequences.

### P*Can*PIE as a near-physiologically active circRNA synthesis platform

Together, these results establish P*Can*PIE as a functional and broadly applicable circRNA synthesis platform. Its fast rate constant and ability to achieve near-complete precursor conversion to circRNA (95 %) at 25 °C distinguish it from all previously described PIE systems, including the *Anabaena* intron (Wesselhoeft et al. 2019) and the recently described CIRC approach (Shen et al. 2025), both of which require 55 °C for efficient circularization. The structural basis for the P*Can*PIE intron temperature advantage may be explained by the lack of an alternative P3 pairing, which has been reported to contribute to kinetic trapping in the *Tetrahymena* intron (Liu et al. 2025; Bonilla et al. 2022). Furthermore, the catalytic core of the *C.a*.mtLSU GpIA1 intron is reinforced by a P7ext-P9.1 tertiary interaction, which is not found in other characterized PIE introns (Liu and Pyle 2024). Together these features bias the intron toward productive folding without requiring thermal rescue.

Although in cellulo activity was not evaluated in this study, the near-physiological splicing conditions of P*Can*PIE suggest it may function in cellular environments, as has been demonstrated for other PIE introns (Ford and Ares 1994; Umekage and Kikuchi 2009). Future directions include testing the P*Can*PIE designs in cellular contexts, incorporating translational elements such as IRES sequences into the circRNA payload for protein expression applications, and exploring strategies for scarless circularization by integrating the spacer sequences into the intron structure itself.

## MATERIALS AND METHODS

### Construct design and cloning

DNA fragments encoding the T7 promoter, permuted intron-exon (PIE) constructs, and terminal BamHI or EcoRI restriction sites were commercially synthesized (IDT, Twist Bioscience) and cloned into the pcDNA3.1, pT7, or pTwist vectors. PIE constructs were designed by repositioning the 5’ and 3’ ends of the C. albicans mtLSU group I intron to peripheral stems P5, P6b, or P8 (Fig. 2B). Short sequence adjustments were introduced at each permutation site to maintain local base pairing. Spacer sequences flanking the circRNA payload were designed to be structurally neutral, replacing the native mtLSU-derived sequences at the junction regions. Spacer lengths of 5, 10, and 20 nt were tested. All construct sequences are provided in Supplementary Table S1.

### In vitro transcription

Plasmid templates were linearized by BamHI or EcoRI (New England Biolabs). Non-radiolabeled RNA was transcribed in reactions containing 1x transcription buffer (400 mM Tris-HCl pH 7.5, 20 mM spermidine, 100 mM NaCl, 0.1% Triton X-100), 3 mM MgCl2, 1 mM each NTP, 10 mM DTT, 6 µg linearized plasmid template, 2.4 U/μL RNaseOUT (ThermoFisher), and 100 µg/mL T7 RNA polymerase (P266L mutant, expressed and purified in-house). Reactions were incubated at 37°C for 2-4 hours.

Radiolabeled RNA for splicing assays was transcribed under the same conditions with the following modifications: UTP was reduced to 1 mM and supplemented with 50 µCi α-32P UTP (3000 Ci/mmol); ATP, GTP, and CTP were each present at 3.6 mM each. Following transcription, radiolabeled precursor RNA was purified by denaturing polyacrylamide gel electrophoresis (see below), excised from the gel, eluted overnight in gel elution buffer (10 mM Na-MOPS pH 6.0, 300 mM NaCl, 1 mM EDTA), recovered by ethanol precipitation, resuspended in RNA storage buffer (6 mM Na-MES pH 6.0) and stored at −80 °C prior to use in splicing assays.

### Non-radiolabeled circRNA synthesis

Following in vitro transcription, reactions were treated with Turbo DNase™ (ThermoFisher) to digest the plasmid template. A 10 µL aliquot was removed as the pre-splicing control. Splicing was initiated in the remaining reaction by addition of MgCl2 to a final supplemental concentration of 0.5 mM and incubated at 25 °C. A 10 µL aliquot was removed following splicing. The remaining material was treated with 0.4 U/μL of RNase R (BioSearch Technologies, RNR07250) to digest linear RNA species and enrich circular RNA products. Pre-splicing, post-splicing, and RNase R-treated samples were resolved by denaturing PAGE or agarose gel electrophoresis for analysis.

### Radiolabeled splicing assays and kinetic analysis

Splicing reactions were assembled using purified radiolabeled precursor RNA. 1 μL of 200 nM precursor RNA, 2 μL of 500 mM HEPES pH 7.5, and 5 μL of 600 mM KCl were mixed and heated to 90 °C for 2 min and cooled at 25 °C for 2 minutes to allow refolding. 3 μL of 30 mM MgCl_2_ was added to a final concentration of 6 mM and the mixture was incubated at 25 °C for 5 minutes. 1 μL of DMSO was added to 5 % (v/v) and the mixture was incubated for a further 5 minutes at 25 °C. Splicing was initiated by addition of guanosine to a final concentration of 40 µM.

1 μL aliquots were removed at 0, 1.5, 3, 5, 7.5, 10, 20, 30, 45, and 60 minutes and immediately quenched by addition of 3x volume of formamide loading buffer (95% formamide, 18 mM EDTA, 0.025% SDS, bromophenol blue, xylene cyanol). Quenched samples were resolved by 0.4 mm-thickness 5 % denaturing polyacrylamide gel and visualized by phosphorimaging. Band intensities corresponding to precursor RNA and circRNA product were quantified using GraphPad Prism.

### RT-PCR and sequencing

CircRNA identity was confirmed by reverse transcription PCR (RT-PCR) using the UltraMarathonRT (uMRT) reverse transcriptase as previously described (Warkentin and Pyle 2025). Briefly, circRNAs were reverse-transcribed using uMRT at ambient temperature with a gene-specific primer that does not overlap with the circRNA ligation junction. PCR was performed using divergent primers flanking the junction to amplify concatemeric products spanning the back-splice junction. RT-PCR amplicons were resolved on a 2% agarose gel in 1x TBE containing GelRed and submitted directly for nanopore sequencing (Plasmidsaurus).

### Denaturing polyacrylamide gel electrophoresis

RNA samples were resolved on 5% polyacrylamide gels containing 7 M urea in 1x TBE buffer. Gels were run at 200 V for 30 minutes. Non-radiolabeled RNA was stained with GelRed (Biotium) and visualized using a ChemiDoc MP (BioRad). Radiolabeled RNA was visualized by phosphorimaging using a Typhoon RGB Biomolecular imager (Cytiva).

### Agarose gel electrophoresis

Non-radiolabeled RNA samples were resolved on 2% agarose gels in 1x TBE buffer containing GelRed. Gels were run at 90 V for 1 hour and imaged under UV illumination.

### Secondary structure prediction

Secondary structure predictions of PIE precursor RNAs and spacer junction sequences were performed using Mfold (Zuker 2003) with default parameters. Predicted structures were used to guide the design of neutral spacer sequences and to identify potential intramolecular interactions between native mtLSU exon sequences and the intron.

### Data Analysis

Band intensities corresponding to precursor RNA and circRNA product were quantified using ImageQuant 8.2 software (GE Healthcare). Band intensity was corrected for the uridine content of each species. The precursor depletion time course data were analyzed and graphed using GraphPad Prism. The fraction of precursor RNA remaining at each time point was plotted as a function of time and fit to a single exponential decay equation:

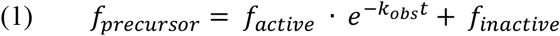

Where *f*_*precursor*_ is the fraction of precursor, *f*_*active*_ is the fraction of actively converting precursor, *k*_*obs*_ is the observed rate, *t* is time (min) and *f*_*inactive*_ is the fraction of non-converting precursor.

All reaction time courses were performed in triplicate and error bars in the kinetic plots were reported as mean ± standard deviation from three independent replicates.

## ACKNOWLEDGMENTS

1) Conceptualization, R.W. and A.M.P.; Investigation, R.W. Writing – Original Draft, R.W.; Writing— Review &Editing, A.M.P.; Funding acquisition, A.M.P.; Supervision, A.M.P. 2) This work was supported by the Howard Hughes Medical Institute (HHMI) for A.M.P. and the Gruber Foundation for R.W. A.M.P is an HHMI investigator.

